# Perilipin 5 interacts with Fatp4 at membrane contact sites to promote lipid droplet-to-mitochondria fatty acid transport

**DOI:** 10.1101/2022.02.03.479028

**Authors:** Gregory E. Miner, Christina M. So, Whitney Edwards, Laura E. Herring, Rosalind A. Coleman, Eric L. Klett, Sarah Cohen

## Abstract

Cells adjust their metabolism by remodeling membrane contact sites that channel metabolites to different fates. Lipid droplet (LD)-mitochondria contacts change in response to fasting, cold exposure, and exercise. However, their function and mechanism of formation have remained controversial. We focused on perilipin 5 (Plin5), an LD protein that tethers mitochondria, to probe the function and regulation of LD-mitochondria contacts. We demonstrate that efficient LD-to-mitochondria fatty acid (FA) trafficking and ß-oxidation during starvation of myoblasts requires both phosphorylation of Plin5 and an intact Plin5 mitochondrial tethering domain. We further identified the acyl-CoA synthetase, Fatp4 (ACSVL4) as a novel mitochondrial interactor of Plin5. The C-terminal domains of Plin5 and Fatp4 constitute a minimal protein interaction capable of inducing organelle contacts. Our work suggests that starvation leads to phosphorylation of Plin5, lipolysis, and subsequent channeling of FAs from LDs to Fatp4 on mitochondria for conversion to fatty-acyl-CoAs and subsequent oxidation.

## Introduction

Cells must adjust their metabolism to respond to changing environmental or developmental cues. An emerging theme is that cells channel metabolites between cellular compartments at sites of close membrane apposition, called membrane contact sites (Prinz, Toulmay and Balla, 2019). These contact sites can be rapidly remodeled in response to signaling cues (Bohnert, 2020). In response to starvation, cytoplasmic lipases release fatty acids (FAs) from neutral lipids stored within the primary fat storage organelle, lipid droplets (LDs) (Zechner, Madeo and Kratky, 2017). FAs are then channeled into mitochondria, where they are oxidized. LD-mitochondria contacts increase in heart and muscle in response to fasting (Wang, Sreenivasan, *et al*., 2011) or exercise (Daemen, Van Polanen and Hesselink, 2018), and in tissue culture cells in response to starvation (Herms *et al*., 2015; Rambold, Cohen and Lippincott-Schwartz, 2015; Nguyen *et al*., 2017; Valm *et al*., 2017). These observations suggest that membrane contact sites may play a role in channeling lipids from LDs to mitochondria for oxidation. However, the function of contacts between LDs and mitochondria remains controversial. For example, in brown adipose tissue, LD-mitochondria contacts decreased during cold exposure. In this situation, LD-associated mitochondria promoted FA storage rather than oxidation (Benador *et al*., 2018). As a result, multiple roles for LD-mitochondria contact sites have been proposed. These roles include: (1) channeling FAs from LDs to mitochondria for oxidation; (2) channeling FAs or FA breakdown products from mitochondria to LDs to protect mitochondria from lipotoxicity; and (3) channeling ATP from mitochondria to LDs to promote neutral lipid synthesis (Wang and Sztalryd, 2011; Benador *et al*., 2019).

Perilipin 5 (Plin5) is one of only several proteins known to mediate LD-mitochondria contact sites (Schuldiner and Bohnert, 2017; Olzmann and Carvalho, 2018). Plin5 is a member of the conserved family of perilipin LD proteins. It is expressed in highly oxidative tissues including heart, skeletal muscle, and brown adipose tissue (Kimmel and Sztalryd, 2016). Superresolution microscopy has localized Plin5 specifically to sites of apposition between LDs and mitochondria (Gemmink *et al*., 2018). Plin5 consists of an N-terminal domain homologous to those of other perilipins, and a unique C-terminal region that recruits mitochondria to LDs (Wang, Sreenivasan, *et al*., 2011). Depletion or overexpression of Plin5 in mouse models has led to apparently contradictory results. Plin5-null mice have fewer LDs in heart and muscle (Kuramoto *et al*., 2012; Mason *et al*., 2014; Drevinge *et al*., 2016; Zheng *et al*., 2017), while overexpression of Plin5 in the heart leads to cardiac steatosis (Wang *et al*., 2013), suggesting a role for Plin5 in fatty acid sequestration. Consistent with these results, cardiomyocytes cultured from Plin5-deficient mice more actively oxidized FAs (Kuramoto *et al*., 2012). In contrast, Plin5 depletion led to reduced FA oxidation in liver (Montgomery *et al*., 2019), while overexpression of Plin5 in skeletal muscle or brown adipose tissue promoted oxidative gene expression (Bosma *et al*., 2013; Gallardo-Montejano *et al*., 2021). Some of these cell type-specific effects may be explained by the idea that phosphorylation of Plin5 by protein kinase A can act as a switch between lipolytic barrier and pro-lipolytic functions of Plin5 (Pollak *et al*., 2015; Keenan *et al*., 2021). Because of these contradictory phenotypes, it has been challenging to untangle the different potential functions of Plin5. In addition, the mitochondrial protein that Plin5 interacts with to mediate membrane contact sites has yet to be identified.

Here, we have carefully dissected the lipolytic barrier and tethering functions of Plin5. By expressing truncated and chimeric protein constructs, we identified a minimal Plin5 C-terminal domain necessary and sufficient to mediate LD-mitochondria contacts. Using fluorescent and radioactive FA pulse-chase assays, we found that both phosphorylation of Plin5 at serine 155 and an intact mitochondrial tethering domain are necessary for efficient channeling of FAs from LDs to mitochondria during starvation of C2C12 myoblasts. We further identified Fatp4 (ACSVL4), an acyl-CoA synthetase (Herrmann *et al*., 2001), as the mitochondrial binding partner of Plin5. Chimeric constructs containing the tether domains of Plin5 or FatP4 were sufficient to create artificial membrane contact sites at the LD-peroxisome or mitochondria-peroxisome interface. Finally, we show that both phosphorylation of Plin5 and Plin5-FatP4 interaction are necessary for efficient channeling of FAs from LDs to mitochondria.

## Results

### Plin5 contains a C-terminal domain sufficient to promote LD-mitochondria contacts

*In vitro* and *in vivo* studies have shown that Plin5 promotes FA storage in LDs and facilitates membrane contact sites between LDs and mitochondria (LD-mito contacts). The N-terminus of Plin5 contains a well characterized lipolytic barrier region that promotes FA storage and is conserved with other perilipins, while the unique C-terminal region is responsible for tethering LDs to mitochondria (Wang, Sreenivasan, *et al*., 2011). Previous studies demonstrated that expression of a chimeric protein fusing the C-terminal region of Plin5 (residues 396-463) to Plin2, a member of the perilipin family, conferred the ability to recruit mitochondria to LDs. Further, it was shown that partial truncation of the C-terminus reduces the ability of Plin5 to induce LD-mito contacts (Wang, Sreenivasan, *et al*., 2011). While the C-terminal region of Plin5 clearly mediates LD-mito contacts, the mitochondrial binding partner of Plin5 has not been identified.

To characterize the C-terminal region of Plin5, we sought to define the specific residues responsible for inducing LD-mito contacts. We first utilized I-TASSER (Roy, Kucukural and Zhang, 2010; Yang and Zhang, 2015; Yang *et al*., 2015), a protein structure prediction algorithm which compares primary sequence with known crystal structures, to predict the protein folding of Plin5. I-TASSER modeling suggested that Plin5 is composed of two separate structures, an N-terminal region homologous with the lipolytic barrier region of the perilipin family, and a C-terminal domain spanning residues 425-463, which we have termed the tether domain (Fig. 1A). The tether domain has no known closely related crystal structure. Therefore, we utilized QUARK (Xu and Zhang, 2012, 2013) *de novo* protein modeling to predict the structure of this domain. QUARK modeling predicts that the tether domain is composed of two α-helices with a hydrophobic pocket (Fig. 1B). Intriguingly, modeling also suggested this hydrophobic pocket may bind lipids, supporting a role in FA trafficking.

**Figure 1.**
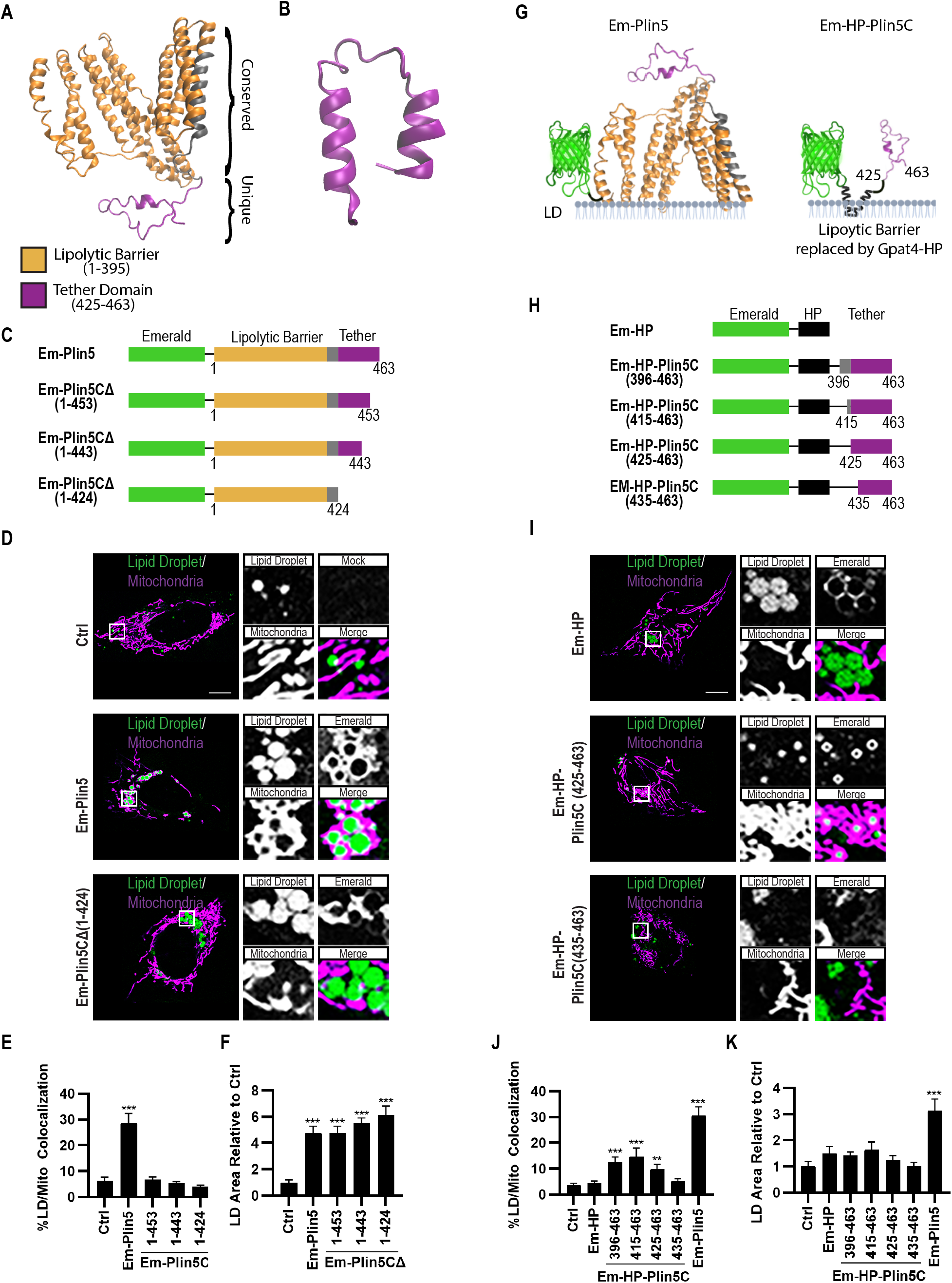
Plin5 contains a C-terminal domain sufficient to promote LD-mito contacts. **(A-B)** Structure predictions of Plin5. (A) I-TASSER structure prediction for Plin5 showing the conserved perilipin lipolytic barrier domain (orange) and a putative C-terminal tether domain with no strong structural prediction (purple). (B) QUARK *de-novo* structure prediction for the putative tether domain shows a two alpha-helix structure containing a hydrophobic pocket. **(C)** Schematic of mEmerald-tagged full-length Plin5 and C-terminal truncations. **(D)** Micrographs of U2OS cells transfected with Mito-RFP and the indicated Em-Plin5 construct, and labeled for LDs with Bodipy 665/676. Merged micrograph includes Mito-RFP and Bodipy 665/667 channels to demonstrate co-localization (white). **(E-F)** Quantification of images from (D). (E) Colocalization of LDs with mitochondria was measured as % LD pixels overlapping with mitochondria, and (F) LD area/cell was measured, normalized to control. **(G-H)** Design of constructs to target the C-terminal region of Plin5 to the LD surface using the hairpin (HP) domain of Gpat4. (G) Cartoon depiction of constructs used to target Plin5 C-terminal region to LD surface. Cartoon created with Biorender.com. (H) Schematic of mEmerald-tagged HP-Plin5 constructs. **(I)** Micrographs of U2OS cells transfected with Mito-RFP and the indicated Em-Plin5 construct, and labeled for LDs with Bodipy 665/676. Merged micrograph includes Mito-RFP and Bodipy 665/667 channels to demonstrate co-localization (white). **(J-K)** Quantification of images from (I). (J) Colocalization of LDs with mitochondria was measured as % LD pixels overlapping with mitochondria, and (K) LD area/cell was measured, normalized to control. Scale bar, 10μm. Data are expressed as means, error bars represent ± SEM. **p < 0.01, ***p < 0.001.

To further define the residues comprising the tether domain, we compared the ability of full-length mEmerald tagged Plin5 (Em-Plin5) to induce LD-mito contacts with a series of Plin5 constructs containing successive C-terminal truncations (Em-Plin5CΔ), guided by our predicted structure (Fig. 1C). As expected, expression of Em-Plin5 in U2OS cells led to a drastic increase in LD-mito colocalization, indicating an increase in membrane contact sites between these organelles. In contrast, cells expressing even a 10-amino acid C-terminal truncation, Plin5CΔ (1-453), exhibited no increase in LD-mito colocalization (Fig. 1D,E). This suggests that both predicted α-helices are essential for mitochondrial tether domain function. We also assessed whether the Plin5 tether domain regulates FA storage. We show that expression of Em-Plin5 and Em-Plin5CΔ constructs led to equivalent increases in LD area (Fig. 1F). These results suggest that Plin5 lipolytic barrier function is independent of LD-mito tethering.

To determine if the tether domain alone was sufficient to induce LD-mito contacts, we created Plin5 constructs lacking the lipolytic barrier region. To maintain LD localization, the lipolytic barrier region of Plin5 was replaced with the hairpin domain (HP) of Gpat4 (residues152-208), a minimal domain that allows targeting to LDs without affecting LD physiology (Wang *et al*., 2016) (Em-HP-Plin5C; Fig. 1G,H). Expression of only the HP domain (Em-HP) demonstrated efficient targeting to the LD, while having no effect on LD-mito contacts (Fig. 1I,J) or LD area (Fig. 1K). Strikingly, expression of Em-HP-Plin5C (396-463) significantly induced LD-mito colocalization. Serial truncations of the C-terminal region demonstrated that expression of the entire tether domain, Em-HP-Plin5C (425-463), is necessary and sufficient to induce LD-mito contacts (Fig. 1J). We also assessed if inducing LD-mito tethering promotes FA storage in the absence of the lipolytic barrier region. Consistent with the truncation experiments shown in Fig. 1F, our results show that expression of the Plin5 tether domain did not affect LD area (Fig. 1K). Together, these results show that removal of even 10 amino acids abrogates Plin5 mitochondrial tether function, but the whole tether domain (425-463) is required to induce LD-mito contacts.

We next tested the functions of the Plin5 lipolytic barrier and tether domains in a more oxidative cell type by expressing a subset of these truncation constructs in C2C12 myoblast cells. As in U2OS cells, we found that the Plin5 lipolytic barrier domain was necessary and sufficient to increase LD area, while the tether domain was necessary and sufficient to increase LD-mito colocalization (Fig. S1A-C). We further examined the relationship between LD size, contacts with mitochondria, and LD movement. LDs are actively transported along microtubules, both at baseline and in response to changing nutritional conditions (Herms *et al*., 2015; Kilwein and Welte, 2019). We hypothesized that increasing LD-mito contacts by overexpressing Plin5 would result in reduced LD motion. Indeed, overexpression of full-length Plin5 significantly slowed LDs in both U2OS (Movies S1-S2) and C2C12 cells (Fig. S1D). Interestingly, expression of either the lipolytic barrier domain (Em-Plin5CΔ) or tether domain (Em-HP-Plin5C) alone also slowed LD motion relative to control, with the tether domain having a larger effect (Movies S3-S4, Fig. S1D). Notably full-length Plin5 slowed LDs to a greater extent than Em-Plin5CΔ or Em-HP-Plin5C.This indicates that both larger LDs and LDs tethered to mitochondria have reduced motion on microtubules, and that Plin5 slows LDs via both mechanisms.

Taken together, our work demonstrates that the Plin5 tether domain, comprised of residues 425-463, is necessary and sufficient to induce LD-mito contacts. Further, our data demonstrates that Plin5 lipolytic barrier activity is independent of LD-mito tethering, but that both lipolytic barrier and tethering domains of Plin5 affect LD motion.

### Plin5 phosphorylation and LD-mito contacts drive ß-oxidation by increasing LD-to-mitochondria FA trafficking

Recent work has suggested that the formation of membrane contact sites, including those between LDs and mitochondria, function to facilitate the rapid transfer of molecules such as lipids. Given that Plin5-induced LD-mito contacts did not promote FA storage, we investigated whether these contacts instead promote trafficking of FAs from LDs to mitochondria under starvation conditions. To assess LD-to-mitochondria FA trafficking, we used an established pulse-chase assay that visualizes FA trafficking using the fluorescent FA analog Bodipy 558/568 C12 (Red C12) (Fig. 2A) (Rambold, Cohen and Lippincott-Schwartz, 2015). C2C12 myoblasts were treated with trace amounts (5 μM) of Red C12 overnight to allow accumulation in LDs, followed by a 1 hr incubation in complete medium (CM) to remove excess label. Red C12 colocalization with LDs and mitochondria was then monitored immediately before (Fig. 2B, CM) and every 30 min following incubation in Hank’s Balanced Salt Solution (HBSS) for 4 hrs. Following incubation in HBSS, cells showed dramatic relocalization of FAs from LDs to mitochondria (Fig. 2B).

**Figure 2.**
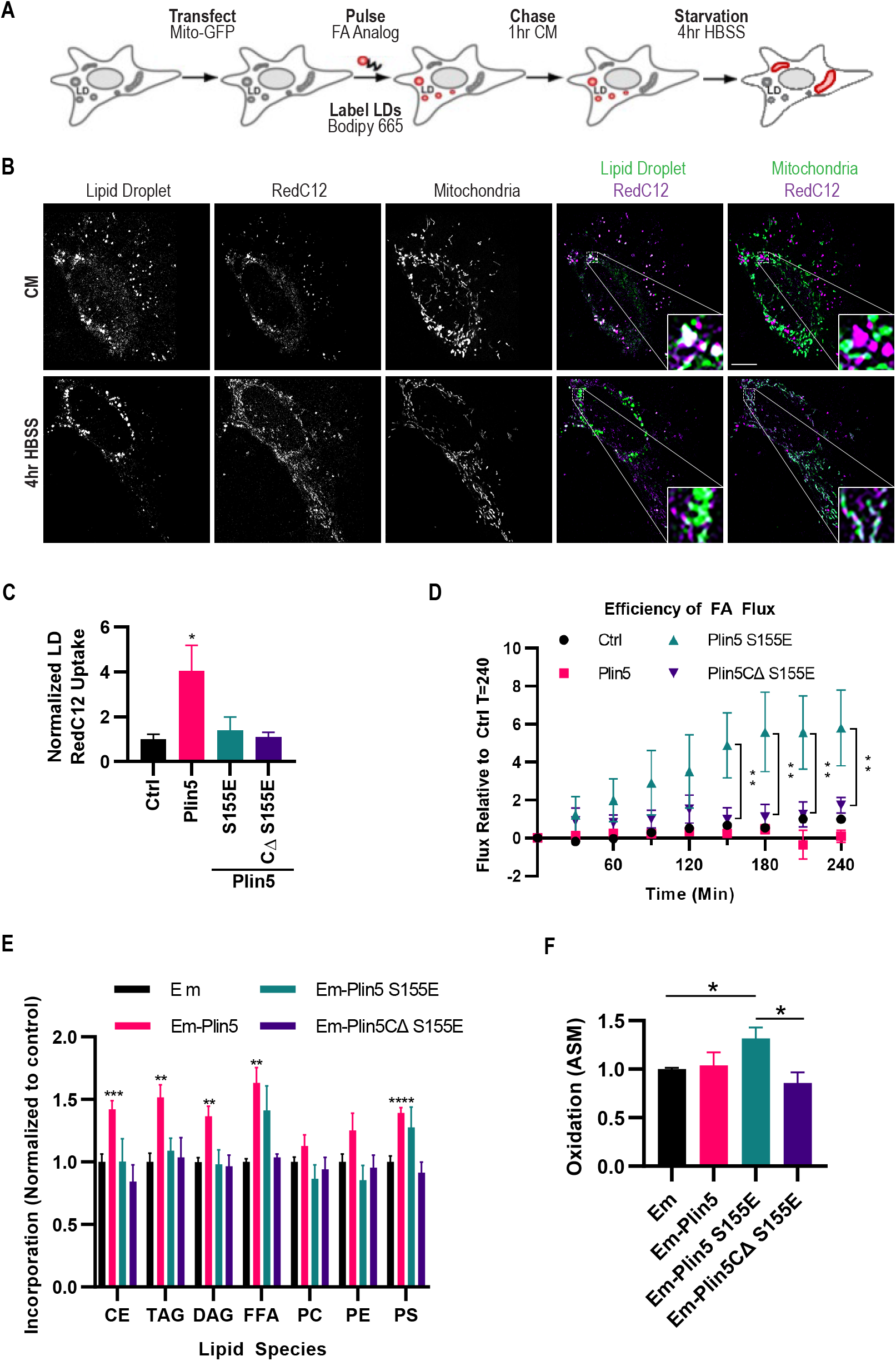
Plin5 phosphorylation and LD-mito contacts drive ß-oxidation by increasing LD-to-mitochondria FA trafficking. **(A)** Schematic representation of FA analog pulse-chase assay design: C2C12 myoblasts were pulsed with FA analog overnight to allow accumulation into LDs, washed, and chased with complete media (CM) for 1 hr. Following chase, incorporation of FA analog into LDs was measured. Cells were then incubated in HBSS for 4 or 6 hrs and FA subcellular localization or oxidation was measured. **(B-D)** Fluorescent pulse-chase assay. Control C2C12 myoblasts or C2C12 myoblasts overexpressing indicated Plin5 constructs were pulsed with Red C12 using the assay described in (A) to monitor FA trafficking from LDs to mitochondria. Following CM chase, cells were imaged to measure incorporation of Red C12 into LDs. Cells were then incubated in HBSS and imaged every 30 min for 4 hrs to monitor FA localization. (B) Representative images of control cells following CM chase or 4 hr HBSS incubation. (C) Red C12 incorporation into LDs following CM chase was measured, normalized to control cells. (D) Red C12 accumulation into mitochondria relative to initial LD incorporation following HBSS starvation was measured, normalized to control cells. **(E-F)** Radioactive pulse-chase assay. Control C2C12 myoblasts or C2C12 myoblasts overexpressing indicated Em-Plin5 constructs were pulsed with a radioactive FA analog [1-^14^C]oleate using the assay described in (A) to monitor FA β-oxidation. Following media chase cells were either collected for TLC to measure incorporation of [1-^14^C]oleate into cellular lipids or incubated in HBSS for 6 hrs to measure β-oxidation. (E) [1-^14^C]oleate incorporation into lipid species following CM chase was measure by TLC, normalized to control cells. Lipid species abbreviations: Cholesterol Ester (CE), Triacylglycerol (TAG), Diacylglycerol (DAG), Free Fatty Acid (FFA), Phosphatidylcholine (PC), Phosphatidylethanolamine (PE), and Phosphatidylserine (PS). (F) β-oxidation following 6 hr incubation in HBSS was assessed by measuring ASM released into media relative to [1-^14^C]oleate incorporation, normalized to control cells. Scale bar, 10 μm. Data are expressed as means, error bars represent ± SEM. *p < 0.05, **p < 0.01, ***p < 0.001 ****p < 0.0001.

Protein kinase A (PKA), a crucial regulator of lipolysis, is activated through phosphorylation during fasting (Pollak *et al*., 2015). Recent studies have shown that Plin5 can also be phosphorylated at Serine 155 by PKA, ablating lipolytic barrier activity and thereby allowing lipolysis (Wang, Bell, *et al*., 2011; Keenan *et al*., 2021). Given that lipolysis is required for LD-to-mitochondria FA trafficking, we created phosphomimetic (S155E) variants of our full length and truncated Plin5 constructs to investigate the contribution of LD-mito contacts to FA trafficking when lipolysis is active. Previous studies found that expression of Plin5 constructs with the same mutation replicated effects of PKA-mediated Plin5 phosphorylation, including changes in cellular localization and transcription (Gallardo-Montejano *et al*., 2016; Najt *et al*., 2020). However, the ability of this phosphomimetic mutation to ablate lipolytic barrier activity has not previously been demonstrated. We first assessed whether expression of our phosphomimetic constructs failed to promote FA storage. As anticipated, cells expressing Plin5 showed increased storage of FAs in LDs in response to a Red C12 pulse; in contrast, cells expressing either Plin5 S155E or Plin5CΔ S155E showed no significant increase in FA storage, indicating that our phosphomimetic mutation inactivated lipolytic barrier activity (Fig. 2C). We attribute decreased FA storage to altered lipolytic barrier activity, as both Plin5 S155E and Plin5CΔ S155E constructs localized appropriately to LDs and affected LD-mito contacts equivalent to Plin5 and Plin5CΔ, respectively (Fig. S2A-B).

We then assessed the ability of our Plin5 constructs to affect FA trafficking during starvation. Cells overexpressing Plin5 failed to traffic FAs from LDs to the mitochondria after 4 hrs in HBSS, suggesting the lipolytic barrier region was not phosphorylated during the time scale of our assay and thus lipolysis could not occur. Strikingly, cells expressing Plin5 S155E showed a dramatic increase in FA trafficking to the mitochondria in response to starvation. In contrast, Plin5CΔ S155E-expressing cells, which have active lipolysis but do not induce LD-mito contacts, showed no significant change in FA trafficking compared to control cells during HBSS incubation (Fig. 2D). These data demonstrate that both Plin5 phosphorylation and LD-mito contacts function to enhance LD-to-mitochondria FA trafficking under starvation conditions.

To investigate whether the observed effects of Plin5 expression on FA trafficking led to changes in FA esterification and β-oxidation, we used a modified version of our pulse-chase assay described above by using trace amounts (17 μM) of a radioactive FA analog, [1-^14^C]oleate. Using thin-layer chromatography (TLC) we confirmed incorporation of [1-^14^C]oleate into LD-specific neutral lipids, including triacylglycerol (TAG), diacylglycerol (DAG) and cholesterol esters (CE), with TAG being the most abundantly labeled species (Fig. 2E). Expression of Plin5 led to increased incorporation of [1-^14^C]oleate into LD-specific neutral lipids, while Plin5 S155E did not (Fig. 2E). This confirms the ability of our phosphomimetic construct to ablate Plin5 lipolytic barrier activity. Intriguingly, we observed that overexpression of Em-Plin5 led to a significant increase in incorporation of [1-^14^C]oleate into phosphatidylserine (PS). Membrane contact sites between ER and mitochondria facilitate PS transport into mitochondria, where PS is subsequently converted to phosphatidylethanolamine (PE) (Kannan *et al*., 2017). By promoting LD-mito contacts, Plin5 overexpression may indirectly reduce ER-mitochondria contact sites and thereby inhibit the transport and subsequent conversion of PS to PE. To assess β-oxidation, we measured the amount of ^14^C acid soluble molecules (ASM) released from cells following nutrient starvation. Mirroring our FA trafficking assay, we found that expression of Em-Plin5 did not change β-oxidation compared to control. In contrast, expression of Em-Plin5 S155E significantly increased β-oxidation relative to control and Em-Plin5. Finally, expression of truncated phosphomimetic Plin5CΔ S155E failed to increase β-oxidation relative to control and showed reduced β-oxidation compared to Plin5 S155E (Fig. 2F).

Previous work has shown that over extended time periods, overexpression of Plin5 S155E can promote transcription of genes that mediate mitochondrial biogenesis and oxidative function, including carnitine palmitoyltransferase I (Cpt1) (Gallardo-Montejano *et al*., 2016). Increased protein expression of Cpt1 has the potential to affect FA transport into mitochondria and β-oxidation. However, on the time scale of our assays (cells transfected for 24 hrs followed by 4 hr starvation), expression of Plin5 S155E did not significantly increase levels of Cpt1 protein (Fig. S2D). Therefore, the effects of expressing Plin5 S155E in our assays can be attributed to protein-protein interactions and LD-mitochondrial tethering, and not to transcriptional effects.

Together, these data demonstrate that Plin5-induced LD-mito contacts enhance the trafficking of FAs from LDs to mitochondria during starvation, leading to increased β-oxidation. This activity is dependent on phosphorylation of the Plin5 lipolytic barrier region, suggesting that Plin5 mediates the metabolic shift from glycolysis to β-oxidation by regulating both lipolysis and FA trafficking.

### The Plin5 tether domain interacts with acyl-CoA synthetase Fatp4

Our studies thus far have established a physiological consequence of LD-mito contacts induced by the tether domain of Plin5. However, the mechanisms by which these contacts form remains unclear. We hypothesized that the tether domain of Plin5 interacts with one or more outer mitochondria membrane (OMM) proteins. To identify protein interactions specific to the tether domain of Plin5 we performed affinity purification-mass spectrometry (AP-MS) with cells expressing either Em-Plin5 or Em-Plin5CΔ. To understand the contribution of the two predicted α-helices (Fig. 1B) to protein binding, we utilized Em-Plin5CΔ constructs lacking either a single helix (Em-Plin5CΔ(1-443)) or both (Em-Plin5CΔ(1-424)). Em-Plin5 and Em-Plin5CΔ protein complexes were affinity purified from transfected U2OS cells using a high-affinity nanobody for GFP (GFP-Trap). As a control, affinity purification was performed using cells transfected with mEmerald alone (Em). Affinity-purified proteins were then analyzed by LC-MS/MS on a QExactive HF mass spectrometer. Two independent biological replicates were analyzed in duplicate. Proteins were considered putative Plin5 interactors if they showed 2-fold enrichment over control and were detected by at least 2 peptides in both biological replicates. Putative interactors were then analyzed using SAINT (Significance Analysis of INTeractome) (Choi *et al*., 2012) to identify Plin5 interactors with high confidence.

A total of 59 putative Plin5 interactors passed our statistical cutoffs (Supplementary Table 1). Of these, 24 showed decreased abundance in both of our affinity-purified Plin5 CΔ protein complexes (Fig. 3A,B) suggesting they interact specifically with the tether domain of Plin5. For most tether-specific interactors (19/24), we found that full truncation of the tether domain led to a 2-fold or greater reduction in protein binding relative to partial truncation, suggesting the α-helix spanning residues 425-443 may be primarily responsible for proteinprotein interactions. We identified two proteins that were of particular interest due to their mitochondrial localization and role in in FA metabolism: Fatp4 and Acot9. Acot9 is primarily localized to the inner mitochondrial membrane (Steensels *et al*., 2020). Therefore, we deemed it unlikely to facilitate organelle tethering, though this data may suggest the formation of a complex spanning mitochondrial membranes. Fatp4 is an OMM bifunctional Acyl-CoA synthetase (ACSVL) that has FA transport and acyl-CoAs synthetase activity, with substrate preference towards C16-C24 FAs (Hall *et al*., 2005). Fatp4 has also been implicated in promoting β-oxidation through re-esterification of long chain FAs following lipolysis (Jia, Moulson, *et al*., 2007; Jia, Pei, *et al*., 2007; Lobo *et al*., 2007). Related ACSL proteins such as ACSL1 have been shown to associate with LD tethering proteins such as Snap23 (Young *et al*., 2018), supporting a potential role in mediating LD-mito contacts. Further, exercise increases expression of both Plin5 (Shepherd *et al*., 2013) and Fatp4 (Jeppesen *et al*., 2012), which may be primarily responsible for the observed increase in LD-mito contacts (Tarnopolsky *et al*., 2007; Devries *et al*., 2013).

**Figure 3.**
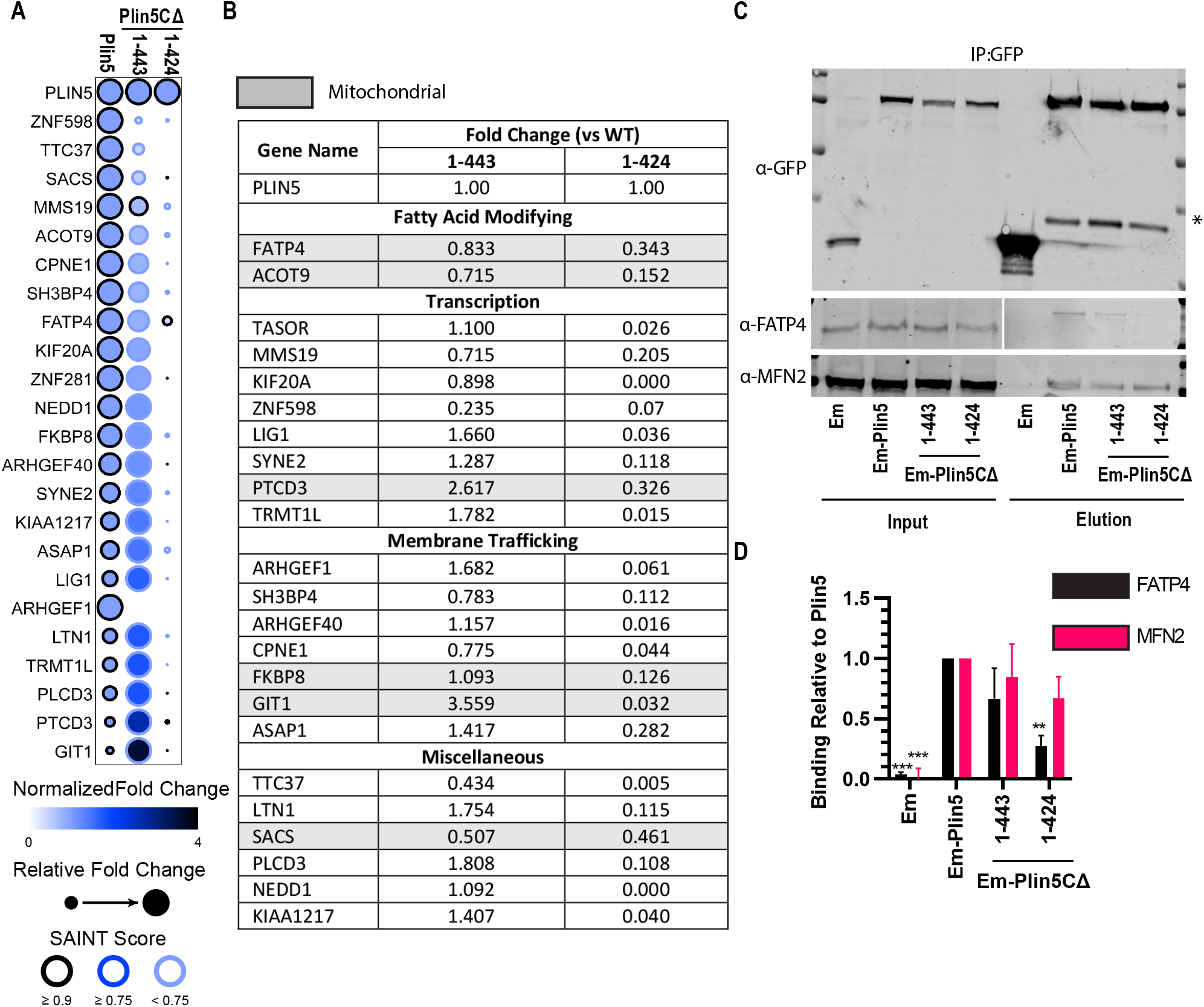
The Plin5 tether domain interacts with acyl-CoA synthetase Fatp4. **(A-D)** Plin5 protein complexes were affinity purified from U2OS cells transfected with the indicated Em-Plin5 construct and analyzed by mass spectrometry. (A) Graphical representation of Plin5 associated protein complexes which show decreased abundance in Plin5CΔ affinity purifications. Circle size and color represent relative enrichment, border color represents statistical significance. (B) Calculated fold change for each Plin5 associated protein and proposed function. Proteins highlighted in grey show mitochondrial subcellular localization. (C) Western blot and (D) quantification of Mfn2 and Fatp4 abundance in affinity purified Plin5 protein complexes. Protein abundance is relative to Plin5, normalized to WT-Plin5 affinity purification. Data are expressed as means, error bars represent ± SEM. *p < 0.05, **p < 0.01, ***p < 0.001.

We confirmed that Fatp4 interacts with the tether domain of Plin5 by western blot analysis of our affinity-purified Plin5 complexes (Fig. 3C). While it did not appear in our AP-MS screen, Mitofusin 2 (Mfn2) has recently been implicated as a potential LD-mito tether through interaction with Plin1 (Boutant *et al*., 2017). Because the lipolytic barrier domains of Plin1 and Plin5 are conserved, we tested whether Plin5 interacts with Mfn2. We show that Mfn2 is in fact a novel Plin5 interactor; however, in contrast to Fatp4, Mfn2 binding to Plin5 does not decrease upon truncation of the tether domain (Fig. 3C,D). This finding is consistent with Mfn2 interacting with both Plin1 and Plin5 via the homologous lipolytic barrier region.

### Plin5 and Fatp4 drive membrane contact site formation

Given that Fatp4 interacts specifically with the Plin5 tether domain, we next investigated whether these proteins constitute a minimal protein interaction sufficient to drive membrane contact site formation. If these proteins constitute a minimal membrane tether they should be capable of inducing contact site formation in the absence of additional LD or mitochondrial proteins. To test this possibility, we first assessed whether recruitment of the Plin5 tether domain to peroxisomes could induce peroxisome-mitochondria contact sites. We adapted our LD-targeted construct design by replacing the HP domain with the peroxisomal membrane protein targeting sequence (residues 1-33) of Pex3 (Pex) to localize the Plin5 tether domain to peroxisomes (Pex-Em-Plin5C) (Fig. 4A-B). Expression of control constructs containing only mEmerald tagged Pex (Pex-Em) in U2OS cells showed efficient recruitment to peroxisomes while having no apparent effect on peroxisome-mitochondria colocalization. In contrast, expression of Pex-Em-Plin5C did cause a significant increase in peroxisome-mitochondria colocalization (Fig. 4C-D). Expression of Pex-Em-Plin5C did not affect peroxisome area relative to control (Fig. S3A). These results suggest that the ability of the Plin5 tether domain to drive membrane contact site formation is not dependent on the unique membrane architecture of the LD.

**Figure 4.**
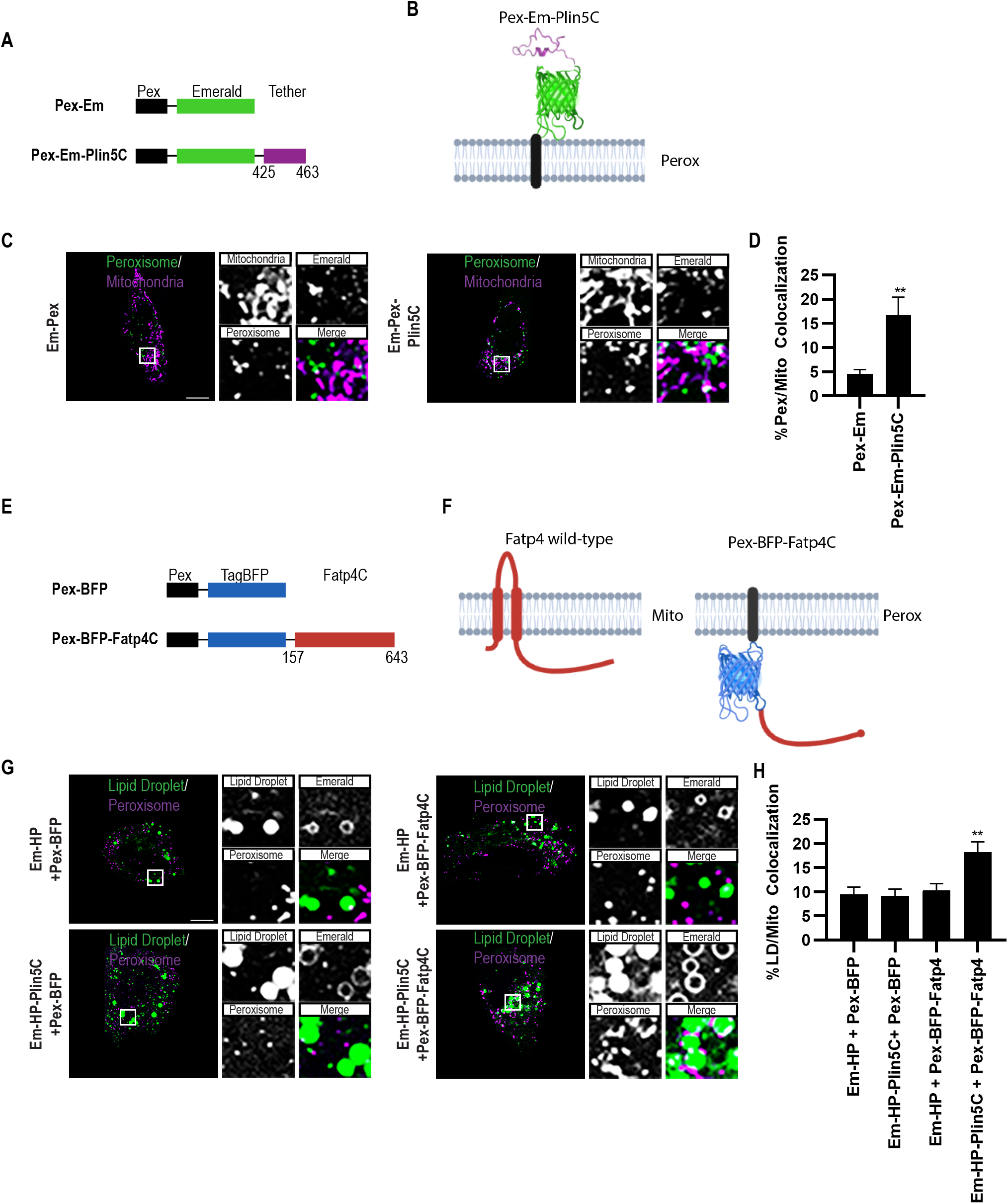
Plin5 and Fatp4 drive ectopic membrane contact sites. **(A-B)** Design of constructs targeting the tether domain of Plin5 to the cytosolic face of the peroxisome membrane using the membrane targeting sequence of Pex3 (Pex). (A) Schematic of mEmerald-tagged Pex constructs. (B) Cartoon depiction of construct used to target Plin5 tether domain to peroxisome membrane. Cartoon created with Biorender.com. **(C)** Micrographs of U2OS cells transfected with mOrange-SKL and indicated Pex construct, mitochondria labeled with MitoTracker Deep Red FM. Merged micrograph includes mOrange-SKL and MitoTracker Deep Red FM channels to demonstrate co-localization (white). **(D)** Quantification of images from (C). Colocalization of peroxisomes with mitochondria was measured as % peroxisome pixels overlapping with mitochondria. **(E-F)** Design of constructs targeting the cytosolic domain of Fatp4 to the cytosolic face of the peroxisome membrane using Pex. (E) Schematic of TagBFP-tagged Pex constructs. (F) Cartoon depiction of construct used to target Fatp4 cytoplasmic domain to peroxisome membrane. **(G)** Micrographs of U2OS cells transfected with mOrange-SKL and indicated HP/Pex targeted constructs, LDs labeled with Bodipy 665/676. Merged micrograph includes mOrange-SKL and Bodipy 665/676 channels to demonstrate co-localization (white). **(H)** Quantification of images from G. Colocalization of LDs with peroxisomes was measured as % LD pixels overlapping with peroxisomes. Scale bar, 10μm. Data are expressed as means, error bars represent ± SEM. **p < 0.01.

We next assessed whether HP-Em-Plin5C can induce contact sites with Fatp4 relocalized to the peroxisomal membrane. Fatp4 is an integral membrane protein containing a large cytosolic C-terminal domain (156-643) (Cho *et al*., 2020), which we hypothesized contained the Plin5 interaction site. For these experiments, we created constructs which localize the C-terminal domain of Fatp4 (Fatp4C) to the peroxisomal membrane (Pex-BFP-Fatp4C) (Fig. 4E,F). Co-expression of Em-HP-Plin5C and Pex-BFP-Fatp4C showed an approximately 2-fold increase in LD-peroxisome colocalization compared to controls transfected with only membrane targeting sequences (Em-HP and Pex-BFP). In contrast, expression of Em-HP-Plin5C or Pex-BFP-Fatp4C alone did not increase LD-peroxisome colocalization (Fig. 4G,H). Expression of these constructs either alone or together did not affect LD or peroxisome area (Fig. S3B,C). Together, these data suggest that the Plin5 tether domain and Fatp4 C-terminal domain form a minimal protein interaction sufficient to drive membrane contact sites. Further, the ability to relocalize either protein while retaining contact site induction suggests additional resident LD or mitochondrial proteins are not required for this activity.

### Loss of Fatp4 impairs Plin5 induced LD-mito contacts and FA trafficking

Recent work to characterize membrane contact sites has shown that many utilize multiple tethers, and loss of a single tether may not drastically alter contact site formation (Prinz, 2014). While we have shown that Plin5 and Fatp4 can interact to induce membrane contact sites, we next tested whether Fatp4 is required for Plin5-driven LD-mito contacts. We assessed the ability of Plin5 to induce LD-mito contacts in cells depleted of Fatp4. Fatp4 expression in C2C12 myoblasts was inhibited by siRNA transfection (Fatp4 siRNA), leading to a >70% reduction in Fatp4 protein abundance relative to non-targeting siRNA (NT siRNA) (Fig. S4A,B). Expression of Em-Plin5 in Fatp4-knockdown cells showed an ~50% reduction in LD-mito contacts relative to NT siRNA controls (Fig. 5A,B). The inability of Fatp4-knockdown to fully ablate Em-Plin5 induction of LD-mito contacts suggests there may be additional Plin5 binding partners on the mitochondrial membrane that facilitate LD-mito contacts. Alternatively, these results may be due to incomplete knockdown of Fatp4 as 10-30% of Fatp4 remained following Fatp4 siRNA knockdown. In contrast, loss of Fatp4 did not appear to affect Plin5 lipolytic barrier function, as expression of Em-Plin5 increased LD area in Fatp4-knockdown cells comparable to NT siRNA treated cells (Fig. 5C).

**Figure 5.**
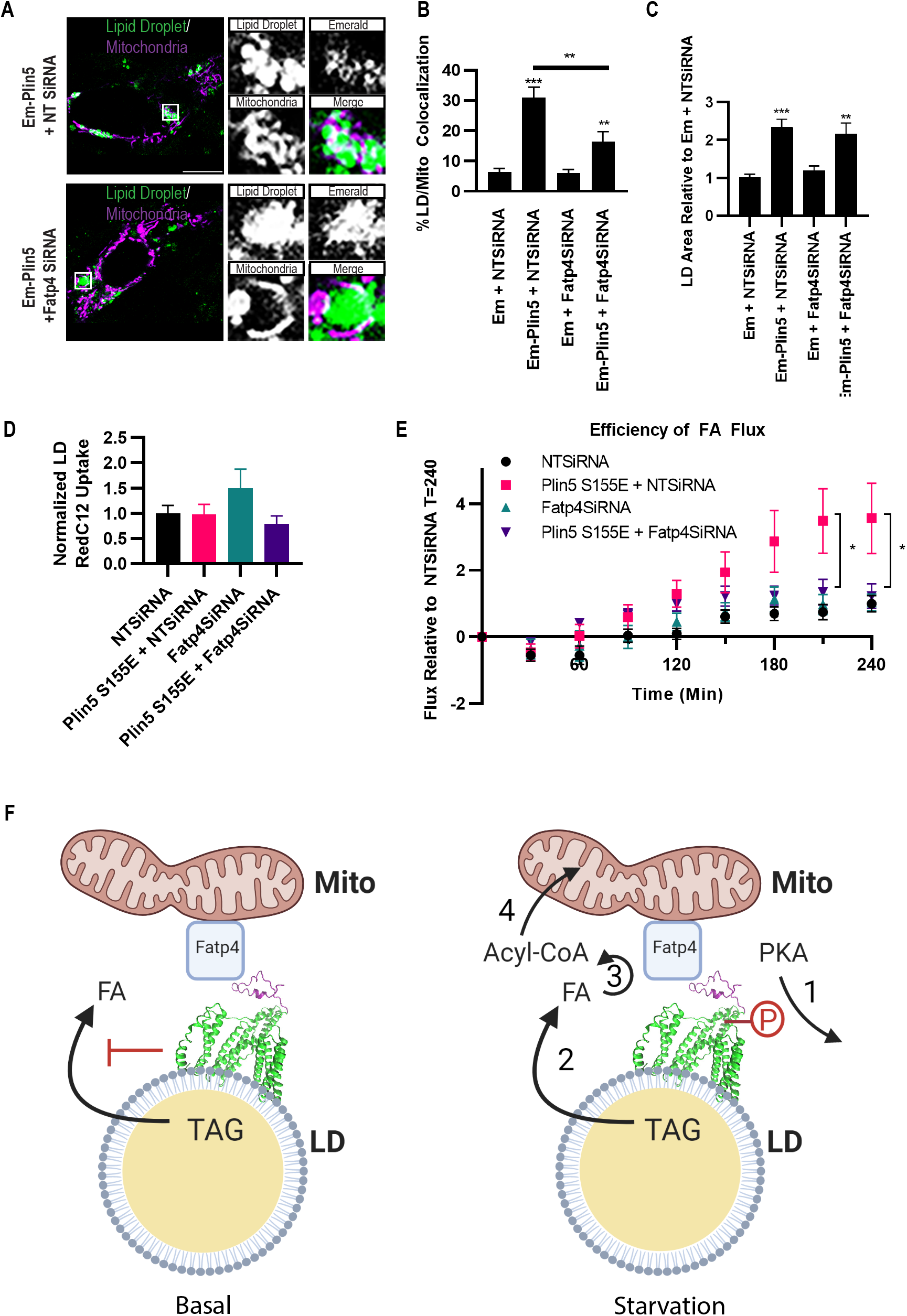
Loss of Fatp4 impairs Plin5-induced LD-mito contacts and FA trafficking. **(A)** Micrographs of C2C12 myoblasts transfected for 24 hr with non-targeting or Fatp4 siRNA. Following RNAi, cells were transfected with Mito-RFP and the indicated Em-Plin5 construct, and LDs labeled with Bodipy 665/676. Merged micrograph includes Mito-RFP and Bodipy 665/676 channels to demonstrate co-localization (white). **(B-C)** Quantification of images from A. (B) Colocalization of LDs with mitochondria was measured as % LD pixels overlapping with mitochondria, and (C) LD area/cell was measured, normalized to control. **(D-E)** C2C12 were assayed as described in Fig. 2A-B, following RNAi treatment using nontargeting or Fatp4 siRNA. (D) Red C12 incorporation into LDs following CM chase was measured, normalized to control cells. (E) Red C12 accumulation into mitochondria relative to initial LD incorporation following HBSS starvation was measured, normalized to control cells. **(F)** Proposed model of FA trafficking during starvation. Under basal conditions Plin5 interacts with ATGL and CGI-58 leading to inhibition of lipolysis. During starvation: (1) PKA phosphorylates Plin5 at S155; (2) phosphorylation of Plin5 promotes lipolysis and release of FAs; (3) released FAs are channeled to mitochondria, where Fatp4 converts FAs to Acyl-CoAs allowing; (4) transport into mitochondria for β-oxidation. Cartoon created with Biorender.com. Scale bar, 10 μm. Data are expressed as means, error bars represent ± SEM. *p < 0.05. **p < 0.01***p < 0.001

Finally, we investigated whether the loss of Fatp4 alters the ability of Plin5 to facilitate LD-to-mitochondria FA trafficking during starvation using the pulse-chase assay shown in Fig. 2A. After Red C12 pulse, Fatp4-knockdown cells transfected with phosphomimetic Plin5 S155E stored a similar amount of RedC12 within LDs compared to NTsiRNA treated cells (Fig. 5D). Upon starvation however, Fatp4-depleted cells expressing Plin5 S155E were unable to efficiently traffic FAs to mitochondria (Fig. 5E). Interestingly, loss of Fatp4 completely ablated Plin5 S155E-induced FA trafficking while only partially preventing LD-mito contacts (Fig. 5B,E). This indicates that Fatp4 not only serves as a LD-mito tether but may also directly facilitate FA trafficking through its ACSL activity. Taken together, our data demonstrate that the tether domain of Plin5 interacts with Fatp4 to induce LD-mito contacts. Further, these Plin5-induced membrane contact sites facilitate trafficking of FAs from LDs to mitochondria during nutrient starvation, promoting β-oxidation.

## Discussion

We have focused on the LD protein Plin5 to investigate the function and mechanism of formation of LD-mito contact sites. We used predictive modeling and expression of truncated and chimeric protein constructs to precisely identify a minimal Plin5 C-terminal domain required for tethering to mitochondria (amino acids 425-463; Fig. 1). The creation of a truncation construct lacking this domain allowed us to identify an acyl-CoA synthetase, Fatp4, as the mitochondrial binding partner of Plin5 by co-immunoprecipitation/mass spectrometry (Fig. 3). Chimeric constructs containing the C-terminal tether domains of Plin5 and Fatp4 were sufficient to create membrane contacts at the LD-peroxisome or mitochondria-peroxisome interface (Fig. 4). We used fluorescent and radioactive FA pulse-chase assays to test the function of LD-mito contacts during starvation of C2C12 myoblast cells. We found that phosphorylation of Plin5 at serine 155, an intact Plin5 mitochondrial tethering domain, and expression of Fatp4 are all necessary for efficient channeling of FAs from LDs to mitochondria and subsequent β-oxidation in response to starvation (Figs. 2 and 5). We propose that during starvation: (1) PKA phosphorylates Plin5 at serine 155 (Keenan *et al*., 2021); (2) phosphorylation of Plin5 promotes lipolysis of triacylglycerol and release of FAs (Wang, Bell, *et al*., 2011; MacPherson *et al*., 2013; Pollak *et al*., 2015); (3) released FAs are channeled from LDs to mitochondria at membrane contact sites, where Fatp4 converts FAs to fatty-acyl-CoAs, allowing; (4) transport into mitochondria for β-oxidation (Fig. 5F).

Our *de novo* modeling of the Plin5 tether domain suggests that this domain contains a hydrophobic pocket that may bind lipids (Fig. 1B). Plin5 has previously been shown to bind lipids, with lipid binding regulated by phosphorylation (Najt *et al*., 2020). FA binding is predicted to be in the Plin5 lipolytic barrier region, though this has not been tested. It is possible that N-terminal lipid binding drives association of Plin5 with LDs, while our identified lipid binding pocket in the tether domain could be for FA transfer to mitochondria. However, whether the Plin5 C-terminal domain has lipid binding and transfer activity remains to be determined.

The presence of acyl-CoA synthetases (ACSLs) at LD-organelle contact sites is an emerging theme. For example, ACSL1 on mitochondria interacts with LD-tethering proteins including SNAP23 (Young *et al*., 2018). Intriguingly, ACSL1 localization at mitochondria appears to be regulated by TANK-Binding Kinase 1 in response to fasting. Relocalization of ACSL1 to mitochondria subsequently increases β-oxidation (Huh *et al*., 2020). Similarly, the ER-LD tethering protein Snx14 interacts with ACSL4 (Datta *et al*., 2020), an isoenzyme of which localizes to ER/LDs (Küch *et al*., 2014). ACSL3 was identified as a protein at the ER membrane microdomains responsible for LD biogenesis, as well as on emerging and mature LDs (Kassan *et al*., 2013), suggesting that ACSL3 may also be present at ER-LD contact sites. The presence of ACSLs at LD-organelle contact sites is consistent with a prominent role of these membrane contact sites in channeling fatty acids to different metabolic fates.

What is the function of Plin5-mediated LD-mito contacts? Multiple functions have been proposed, based on seemingly contradictory results from mouse and cell models of Plin5 ablation or overexpression. Our results are consistent with previous work showing that Plin5 can switch between lipolytic barrier and pro-lipolytic activities, depending on its phosphorylation state (Pollak *et al*., 2015; Keenan *et al*., 2021). Our findings are also consistent with recently published work showing that expression of truncated Plin5 in human AC16 cardiomyocytes does not alter β-oxidation relative to full-length Plin5 during acute starvation (Kien *et al*., 2022). Because that study did not use phosphomimetic Plin5 constructs, the authors concluded that the Plin5 mitochondrial tether domain is dispensable for FA oxidation. Here we have used a combination of phosphomimetic and truncation constructs to carefully tease apart the roles of the Plin5 lipolytic barrier and mitochondria tether domains. While the Plin5 lipolytic barrier domain can promote either FA storage or mobilization depending on its phosphorylation state, we found that the Plin5 mitochondrial tether domain was required for efficient flux of FAs from LDs to mitochondria, and their subsequent oxidation. Thus, we propose that a primary function of Plin5-mediated LD-mito contacts is to efficiently channel FAs from LDs to mitochondria. This occurs through interaction of the Plin5 C-terminal domain with Fatp4. Interestingly, LD-to-mitochondria FA flux was completely ablated by knockdown of Fatp4, while LD-mito colocalization was reduced but not abolished by Fatp4 depletion (Fig. 5). This implies that additional mitochondrial proteins may interact with Plin5 at membrane contact sites, but that the Fatp4 interaction is specifically required for FA channeling. Indeed, we observed an interaction between the Plin5 lipolytic barrier domain and the mitofusin Mfn2 (Fig. 3). Mfn2 was previously reported to interact with Plin1, which shares a conserved N-terminal domain with Plin5 (Boutant *et al*., 2017). Thus, Plin5-mediated LD-mito contacts may have additional functions beyond FA channeling, including the regulation of mitochondrial fusion. Whether the interaction of Plin5 with binding partners including Fatp4 and Mfn2 occurs at the same or distinct membrane microdomains remains to be determined. The spatial relationship between Plin5/Fatp4 and other LD-mito contact proteins including SNAP23/ACSL1 (Jägerström *et al*., 2009; Young *et al*., 2018), MIGA2 (Freyre *et al*., 2019), and VPS13D/TSG101 (Wang *et al*., 2021) is also an area of future investigation.

The channeling of FAs directly from LDs to mitochondria at membrane contact sites during starvation has the benefit of preventing a build-up of cytoplasmic FAs, which can lead to mitochondrial toxicity (Nguyen *et al*., 2017). Alternative FA trafficking routes from LDs to mitochondria have previously been proposed. In hepatocytes, it was suggested that the primary route is through lipophagy of LDs, followed by fusion of lysosomes with the plasma membrane and subsequent reuptake of FAs and transport to mitochondria (Cui *et al*., 2020). Like FA transfer at LD-mito contact sites, lysosomal fusion with the plasma membrane would also prevent a buildup of cytoplasmic FAs, by releasing FAs extracellularly and allowing cytoplasmic FA concentration to be regulated at the reuptake step. Which mechanism is used in a given situation likely depends on the cell-type specific expression of regulatory proteins, as well as the stimulus initiating lipolysis/lipophagy.

In summary, we have identified the acyl-Co-A synthetase Fatp4 as a novel interactor of Plin5. We provide evidence that Plin5 interacts with Fatp4 at LD-mito contact sites to promote LD-to-mitochondria trafficking of FAs in a phosphorylation-dependent manner. This work helps to reconcile seemingly contradictory results seen in models of Plin5 ablation/overexpression by revealing the specific functions of the lipolytic barrier vs. mitochondrial tethering domains of Plin5.

## Acknowledgements

We thank Amit Joshi and current members of the Cohen lab for helpful discussions and critical reading of the manuscript. We are grateful to Reginald Edwards for performing pilot LD tracking experiments. This research is based in part upon work conducted using the UNC Proteomics Core Facility, which is supported in part by P30 CA016086 Cancer Center Core Support Grant to the UNC Lineberger Comprehensive Cancer Center. Research reported in this publication was supported by the National Institute of General Medical Sciences of the National Institutes of Health under award numbers R35 GM133460 (S.C.) and F32 GM136027 (G.E.M.).

## Author Contributions

G.E.M. and S.C. designed the research; G.E.M., C.M.S., W.E., and L.E.H. performed research; G.E.M., C.M.S., W.E., L.E.H., and E.L.K. analyzed data; L.E.H., E.L.K., and R.A.C. provided expertise and feedback, and G.E.M. and S.C. wrote the paper, with input from all co-authors.

## Methods

### Cell Culture and Transfection

U2OS cells and C2C12 myobolasts were obtained from the UNC Tissue Culture Facility and maintained in Dulbecco’s Modified Eagle Medium (DMEM) with 10% FBS and 4 mM glutamine (complete medium, CM). Cells were cultured on chambered cover glass (#1.5 high performance cover glass, Cellvis), coated with 10 μg/ml fibronectin (MilliporeSigma). For starvation experiments, Hank’s buffered salt solution (HBSS, 14025092) was purchased from Thermo Fisher. Cells were transfected with Mito-RFP, Mito-GFP, mOrange-SKL, or Plin5 constructs using Lipofectamine 2000 (Invitrogen) according to the manufacturer’s instructions. RNA interference was performed using 25 nM ON-TARGETplus Mouse Fatp4 (J-063631-12-0002) siRNA or ON-TARGETplus non-targeting pool (Horizon Discovery) and DharmaFECT 1 (Horizon Discovery) according to the manufacturer’s instructions.

### Plasmids

Plasmids used are listed in Supplemental Table 2. Full-length Plin5, Plin5 truncations and Chimeric Plin5 or Fatp4 constructs were generated using HiFi DNA Assembly Master mix (New England Biolabs, E2621). Plin5 mutant constructs were generated using Q5® Site-Directed Mutagenesis Kit (New England Biolabs, E0554S) mEmerald-C1 and BFP-KDEL were used as vectors for mEmerald- and TagBFP-tagged constructs, respectively. For plasmid construction, all PCRs were performed using Q5 High Fidelity DNA polymerase (M0419; New England Biolabs) and restriction enzymes were from New England Biolabs. The following plasmids were kind gifts: pEYFP-C1-Plin5 from Carole Sztalryd (University of Maryland); TagBFP-KDEL from Gia Voeltz (University of Colorado); mEmerald-C1 and mOrange-SKL from Michael Davidson (Florida State University); Mito-GFP and Mito-RFP from Jennifer Lippincott-Schwartz (Janelia Research Campus).

### Inhibitors and Antibodies

The following chemicals, dyes and antibodies were used: 10 μg/ml fibronectin (MilliporeSigma), 5 μM BODIPY 558/568 C12 (Life Technologies), 5 nM [1-^14^C]oleate (Perkin Elmer), 50 ng/ml BODIPY 665/676 (Life Technologies), 100 nM MitoTracker Deep Red (Life Technologies), L-carnitine (C0158-1G, Thermo Fisher), bovine serum albumin-fatty acid free (MilliporeSigma), anti-CPT1B (Abcam, ab104662), anti-GFP (Invitrogen, A10262), anti-Fatp4 (Abcam, ab200353), anti-Mfn2 (Cell Signaling, 11925), anti-BIII-tubulin (Abcam, ab18207), and anti-Plin5 (American Research Products, 03-GP31).

### Fluorescent FA Analog Pulse-Chase

C2C12 cells were incubated with CM containing 5 μM BODIPY 558/568 C12 (Red C12, Life Technologies) for 16 hr. Cells were then washed three times with CM, and chased for 1 hr in order to allow the fluorescent lipids to incorporate into LDs. Cells were imaged immediately following pulse-chase labeling with Red C12. Cells were then washed three times with HBSS and imaged every 30 min for 4 hrs in order to track the subcellular localization of FAs. Cells were transfected with Mito-GFP 24 hrs prior to Red C12 labeling to visualize mitochondria. To label LDs, Bodipy 665/676 (Life Technologies) was added to cells at 50 ng/ml 16 hrs prior to imaging and was present during imaging.

### Radioactive FA Pulse-Chase

C2C12 cells were incubated with Krebs-Ringer bicarbonate HEPES buffer (KRBH) labeling buffer (KRBH containing 5.5 mM glucose, 1mM carnitine, 0.25% FA free BSA) and 17 μM [1-^14^C]oleate (^14^C-oleate, Perkin Elmer) for 16 hrs. Cells were then washed three times with KRBH, and chased for 1 hr in KRBH labeling buffer in order to allow ^14^C-oleate incorporation into cellular lipids. Following the pulse-chase cells were either collected to measure ^14^C-oleate incorporation, or incubated in HBSS to assess β-oxidation.

### Lipid Extraction and TLC

Following the Radioactive FA pulse-chase, cells were washed twice with KRBH containing 1% FA Free BSA. Lipids were isolated by chloroform-methanol extraction. Following extraction lipids were dried by speedvac and resuspended in chloroform-methanol. Aliquoted samples were counted for total ^14^C-oleate incorporation. Remaining samples were spotted onto silica gel preparative TLC plates (MilliporeSigma). Separation of lipids was performed by developing the plates in a 2-step solvent system of chloroform:methanol:ammonium hydroxide, 65:25:4 (vol/vol) followed by heptane:isopropyl ether:acetic acid (15:10:1) (vol/vol). Radiolabeled were visualized using a Bioscan AR-2000 radio-TLC imaging scanner (Bioscan).

### ASM Measurement

Following Radioactive FA pulse-chase, cells were washed three times with HBSS and incubated with HBSS containing 1 mM carnitine. 500 μl of media was collected after 6 hrs and incubated with 12.5 μl of 25% FA-free BSA overnight at 4 °C followed by centrifugation at 20,800 gfor 20 min. Supernatants were washed once more with 12.5 μl of 25% FA-free BSA and spun 20,800 g for 20 min, before aliquots of the supernatant were counted for ^14^C-labeled ASM.

### Isolation of GFP-Plin5 Complexes for Mass Spectrometry

mEmerald protein complex isolation for mass spectrometry was performed as previously described(Kaltenbrun *et al*., 2013)utilizing GFP-Trap magnetic beads (Chromotek). Briefly, U2OS cells expressing mEmerald constructs were washed twice with cold PBS (Thermo Fisher, #14190144), harvested by scraping with a cell lifter, and centrifuged at 350 g for 10 min at 4°C. Cell pellets were resuspended in 20 mL resuspension buffer (20mM HEPES, pH 7.4, 1.2% polyvinylpyrrolidone) with protease (MiliporeSigma, #8340) and phosphatase inhibitors (MiliporeSigma, #5726 and #P0044), and snap frozen in liquid nitrogen. Cells were lysed by cryogenic grinding using a MM 301 Mixer Mill (10 cycles, 2.5 min at 30Hz) (Retsch). Lysate was resuspended in MS lysis buffer (20mM K-HEPES pH 7.4, 150mM NaCl, 100mM KOAc, 2mM MgCl2, 0.1% Tween-20, 1μm ZnCl2 1 μmCaCl2, 0.5% Triton X-100) with protease and phosphatase inhibitors (5 ml/g cells). Resuspended lysate was homogenized using a Polytron (Kinematica) (2 × 15 sec at 25,000 RPM) and pelleted at 2500 g at 4°C. Cleared lysate was incubated with 50 μl equilibrated GFP-Trap magnetic beads by rotating for 1 hr at 4°C. Beads were washed 6 times with 1 ml of lysis buffer. GFP complexes were eluted from beads in 40 μl 1X LDS Sample Buffer (Invitrogen) at 70°C for 15 min. Eluted complexes were alkylated with 100 mM iodoacetamide for 1 hr at room temperature prior to mass spectrometry analysis.

### Liquid Chromatography-Tandem Mass Spectrometry (LC-MS/MS) Analysis

Immunoprecipitated samples were subjected to SDS-PAGE and stained with coomassie. Lanes (1cm) for each sample were excised and the proteins were reduced with 5mM DTT for 30 min at 55C, alkylated with 15mM IAA for 45 min in the dark at room temperature, and in-gel digested with trypsin overnight at 37°C. Peptides were extracted, desalted with C18 spin columns (Pierce) and dried via vacuum centrifugation. Peptide samples were stored at −80°C until further analysis.

Each sample was analyzed in duplicate by LC-MS/MS using an Easy nLC 1200 coupled to a QExactive HF (Thermo Scientific). Samples were injected onto an Easy Spray PepMap C18 column (75 μm id × 25 cm, 2 μm particle size) (Thermo Scientific) and separated over a 60 min method. The gradient for separation consisted of 5–45% mobile phase B at a 250 nl/min flow rate, where mobile phase A was 0.1% formic acid in water and mobile phase B consisted of 0.1% formic acid in 80% ACN. The QExactive HF was operated in data-dependent mode where the 15 most intense precursors were selected for subsequent HCD fragmentation. Resolution for the precursor scan (m/z 350–1600) was set to 60,000, while MS/MS scans resolution was set to 15,000. The normalized collision energy was set to 27% for HCD. Peptide match was set to preferred, and precursors with unknown charge or a charge state of 1 and ≥ 7 were excluded.

Raw data files were processed using Proteome Discoverer version 2.1 (Thermo Scientific). Peak lists were searched against a reviewed Uniprot human database (containing 20,414 protein sequences, downloaded January 2019). appended with a common contaminants database. The following parameters were used to identify tryptic peptides for protein identification: 10 ppm precursor ion mass tolerance; 0.02 Da product ion mass tolerance; up to two missed trypsin cleavage sites; carbamidomethylation of Cys was set as a fixed modification; oxidation of was set as a variable modification. Scaffold (version 4.7.3, Proteome Software) was used to validate MS/MS based peptide and protein identifications, and to provide relative quantitation. Peptide identifications were accepted if they could be established at greater than 95% probability to achieve an FDR less than 0.1% by the Scaffold Local FDR algorithm. Protein identifications were accepted if they could be established at greater than 99.0% probability and contained at least 2 identified peptides. SAINT (Significance Analysis of INTeractome) (Choi *et al*., 2012) was used to identify Plin5 interactors. Proteins were identified as Plin5 interactors if they had a FC_A (Average enrichment relative to Em control) score greater than 2.0 and a SAINT score greater than 0.75. Relative quantitation was performed using the calculated quantitative values (normalized peak area) within Scaffold.

Isolation of GFP-Plin5 Complexes for Co-Immunoprecipitation/Western Blot Isolation of mEmerald protein complexes for co-immunoprecipitation was performed utilizing the same harvesting method used for mass spectrometry. Following cell harvesting, Co-immunoprecipitation was performed utilizing GFP-Trap magnetic beads (Chromotek) following manufacturer protocols with slight modification. Cell pellets were resuspended in 400 μL of Co-IP Lysis Buffer (10 mM Tris/Cl pH 7.5; 150 mM NaCl; 0.5 mM EDTA; 0.5% NP-40) with protease and phosphatase inhibitors. Cell suspension was incubated on ice for 30 min with pipetting every 10 minutes and pelleted at 20,000 g for 10 min at 4 °C. 600 μl of Co-IP Dilution Buffer (10 mM Tris-Cl pH 7.5; 150 mM NaCl; 0.5 mM EDTA) was added to cleared lysate. Diluted lysate was incubated with 25 μl equilibrated GFP-Trap magnetic beads by rotating for 1 hr at 4 °C. Beads were washed 3 times with 500 ul of Co-IP Dilution Buffer. GFP complexes were eluted from beads in 30 μl 6X Laemmli Sample Buffer at 95 °C for 10 min. Protein complexes were examined by Western blotting.

### Microscopy, Image Processing, Analysis, and Statistics

Images were acquired on an inverted Zeiss 800/Airyscan laser scanning confocal microscope equipped with 405, 488, 561 and 647 nm diode lasers, and Galium Arsenide Phosphid (GaAsP) and Airyscan detectors. Confocal and Airyscan images were acquired using a 63x/1.4 NA objective lens, at 37 °C and 5% CO_2_ (Carl Zeiss, Oberkochen, Germany). For movies S1-S4, Airyscan timelapse images were acquired every 27 s for 10 frames. Airyscan images were processed in Zen software (Carl Zeiss) using a processing strength of 6.0.

Image brightness and contrast were adjusted in Adobe Photoshop CS. Images were analyzed using ImageJ (NIH).

For organelle colocalization, masks of each organelle were created using the corresponding channels. For overlap of LDs with organelles (mitochondria, peroxisomes), the percentage of LD pixels colocalized with the organelle versus total LD pixels was calculated. For overlap of peroxisomes and mitochondria the percentage of peroxisome pixels colocalized with the organelle versus total peroxisome pixels was calculated.

LDs were tracked using the TrackMate (Tinevez *et al*., 2017) plugin, an ImageJ implementation of the linear assignment problem (LAP) tracking algorithm (Jaqaman *et al*., 2008).

For fluorescent FA pulse-chase assays, images were analyzed using ImageJ to determine total fluorescence intensity. To quantify the fluorescence intensity of Red C12 in mitochondria and LDs, we made mitochondria and LD masks using the Mito-GFP and Bodipy 665/667 channels respectively, and then the fluorescence intensity in the Red C12 channel was calculated across the entire mask.

Data were expressed as means, error bars represent ± SEM. Statistical analysis among groups was performed using Student’s t test.

## Supplemental Figure Legends

**Supplemental Figure 1.**
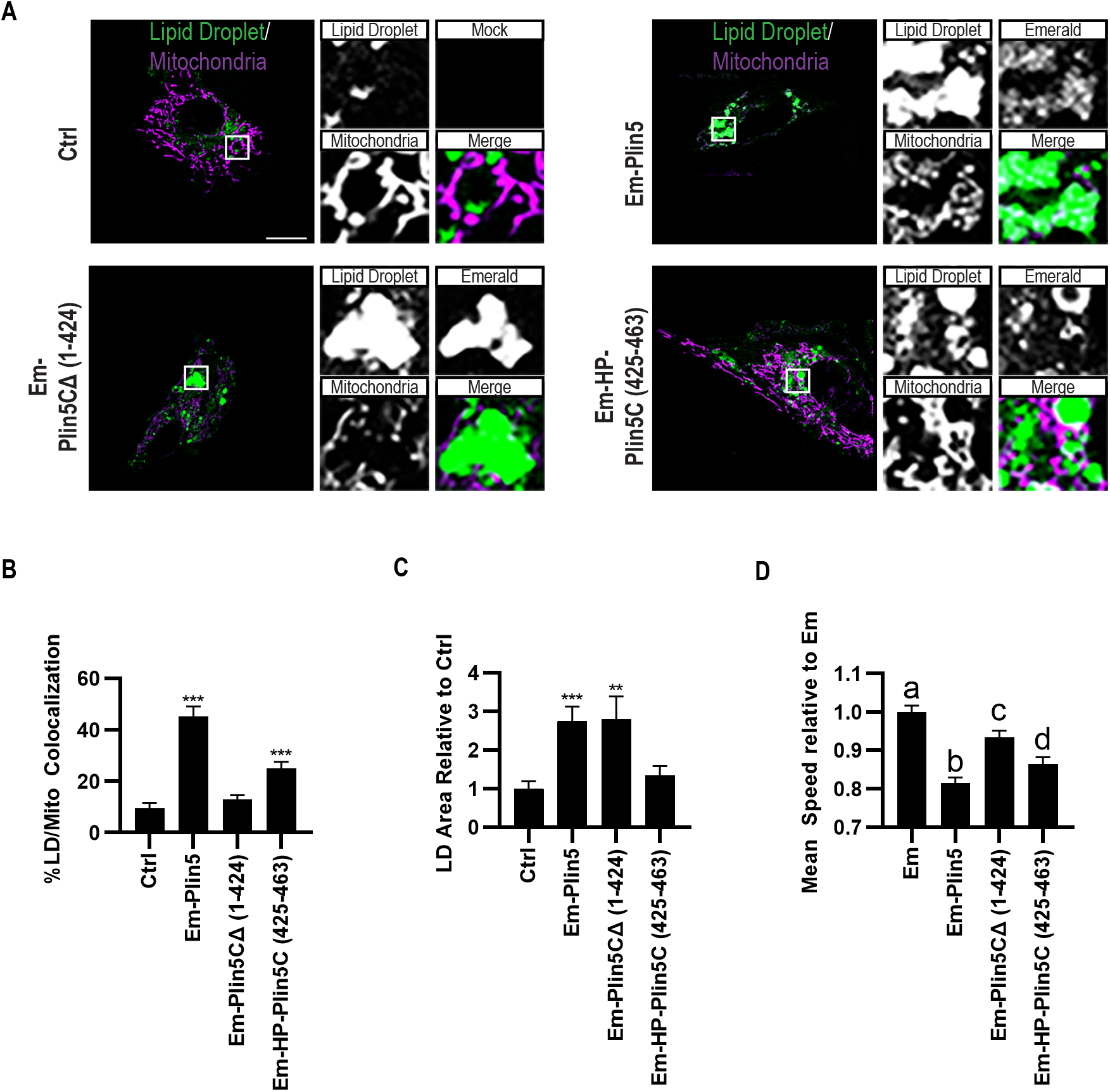
Effect of Plin5 on LD-mito contacts and LD size in C2C12 myoblasts. **(A)** Micrographs of C2C12 myoblasts transfected with Mito-RFP and the indicated Em-Plin5 construct, and labeled for LDs with Bodipy 665/676. Merged micrograph includes Mito-RFP and Bodipy 665/667 channels to demonstrate co-localization (white). **(B-C)** Quantification of images from (A). (B) Colocalization of LDs with mitochondria was measured as % LD pixels overlapping with mitochondria, and (C) LD area/cell was measured, normalized to control. **(D)** Quantification of LD mean speed in C2C12 myoblasts transfected with the indicated Em-Plin5 construct, normalized to myoblasts transfected with Em control. Scale bar, 10 μm. Data are expressed as means, error bars represent ± SEM. **p < 0.01, ***p < 0.001.

**Supplemental Figure 2.**
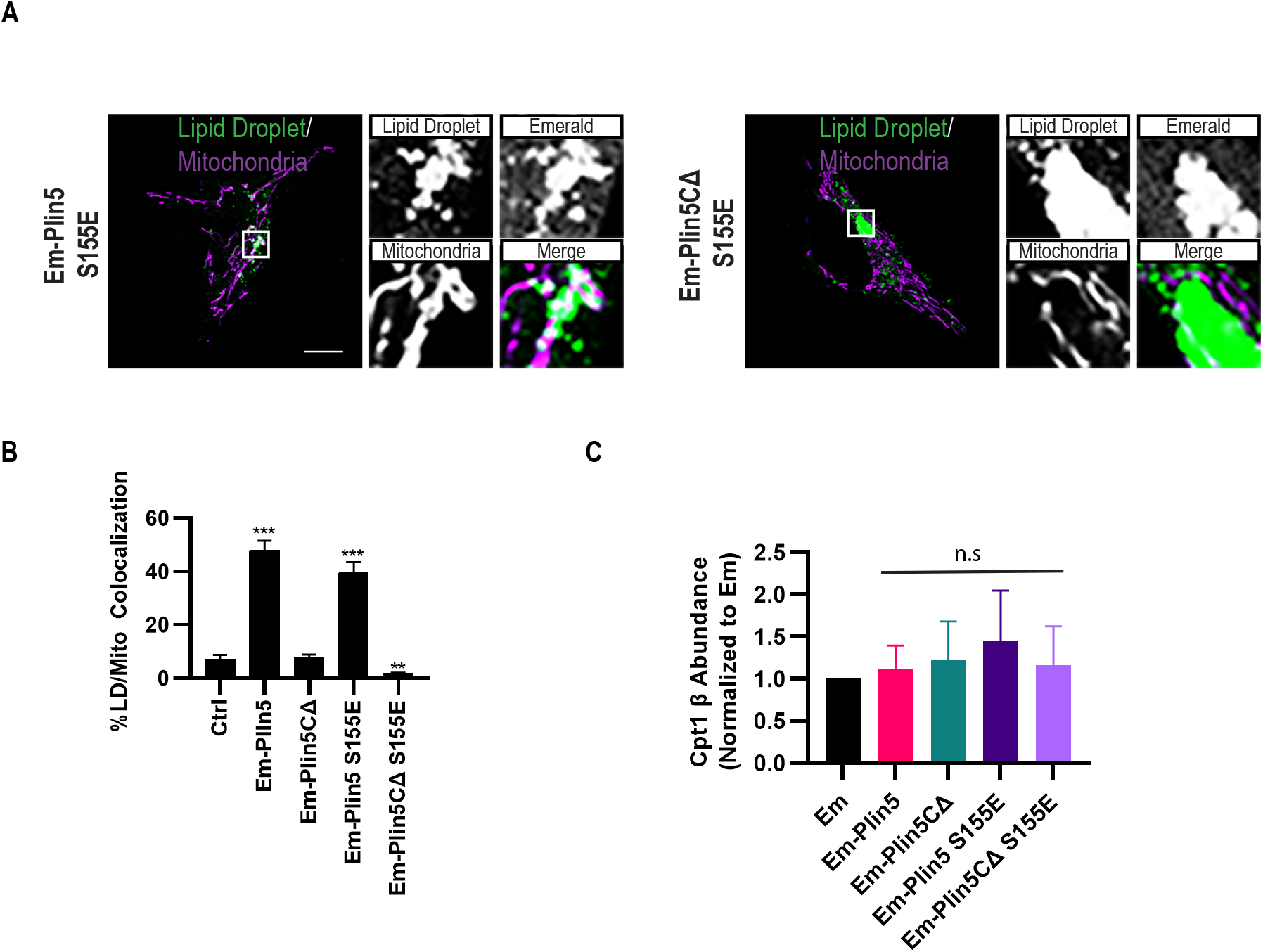
Plin5 phosphomimetic constructs promote LD-mito contacts and do not alter Cpt1b expression in C2C12 myoblasts. **(A)** Micrographs of C2C12 myoblasts transfected with Mito-RFP and indicated Em-Plin5 construct, and labeled for LDs with Bodipy 665/676. Merged micrograph includes Mito-RFP and Bodipy 665/667 channels to demonstrate co-localization as shown by white pixels. **(B)** Quantification of images from A. (B) Colocalization of LDs with mitochondria was measured as % LD pixels overlapping with mitochondria. **(C)** Quantitation of Cpt1b expression in C2C12 myoblasts 48 hrs after transfection with the indicated Em-Plin5 construct, normalized to myoblasts transfected with Em control. Scale bar, 10 μm. Data are expressed as means ± SEM. **p < 0.01, ***p < 0.001

**Supplemental Figure 3.**
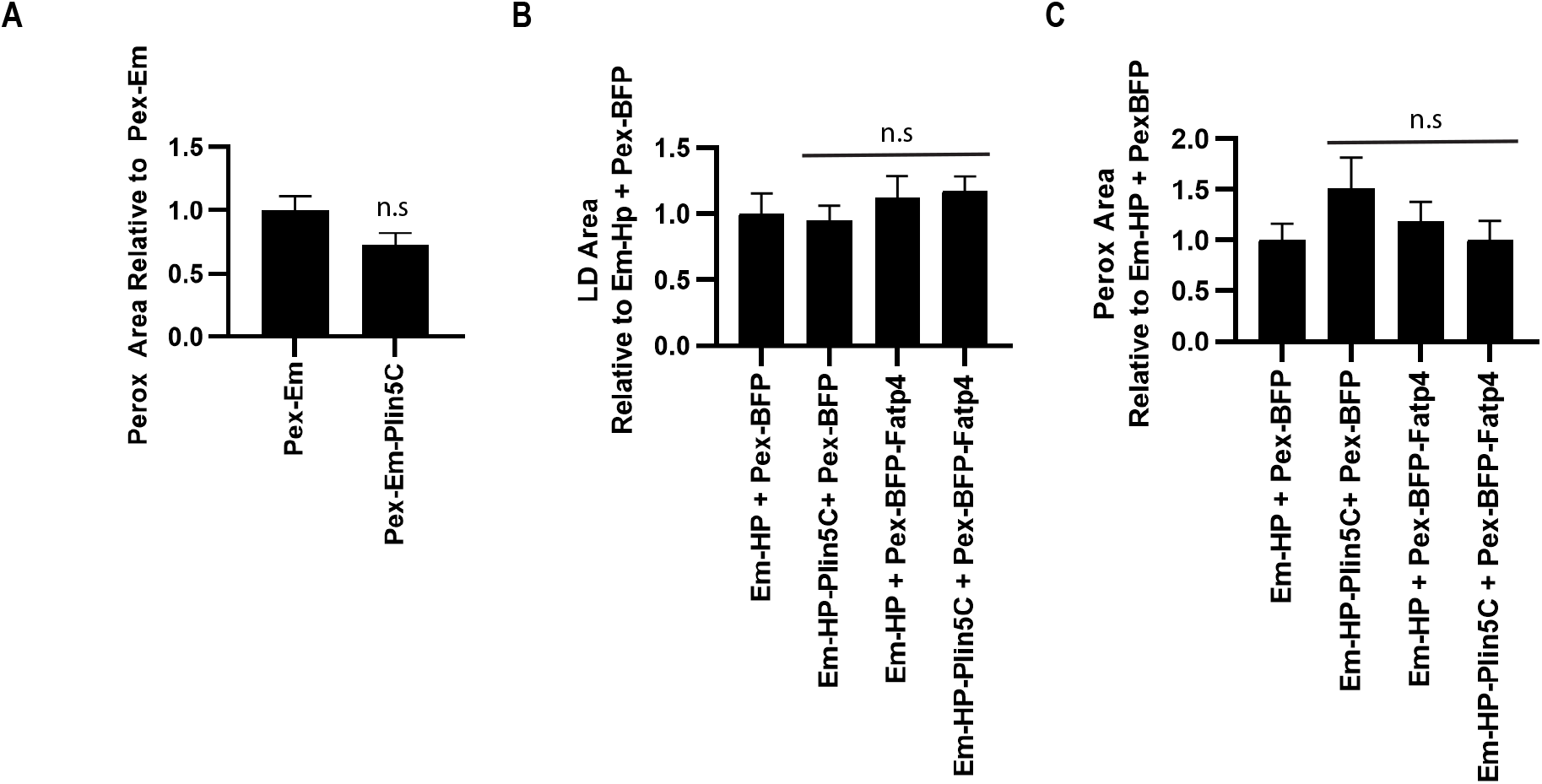
Plin5 and Fatp4 Pex constructs do not alter peroxisome area. **(A-C)** Effect of organelle-targeted constructs on organelle area. (A) Quantitation of the effect of Em-Pex constructs on peroxisome area/cell, normalized to control. (B) Quantitation of the effect of Em-HP and TagBFP-Pex constructs on (B) LD and (C) peroxisome area/cell was measured, normalized to control. Data are expressed as means, error bars represent ± SEM.

**Supplemental Figure 4.**
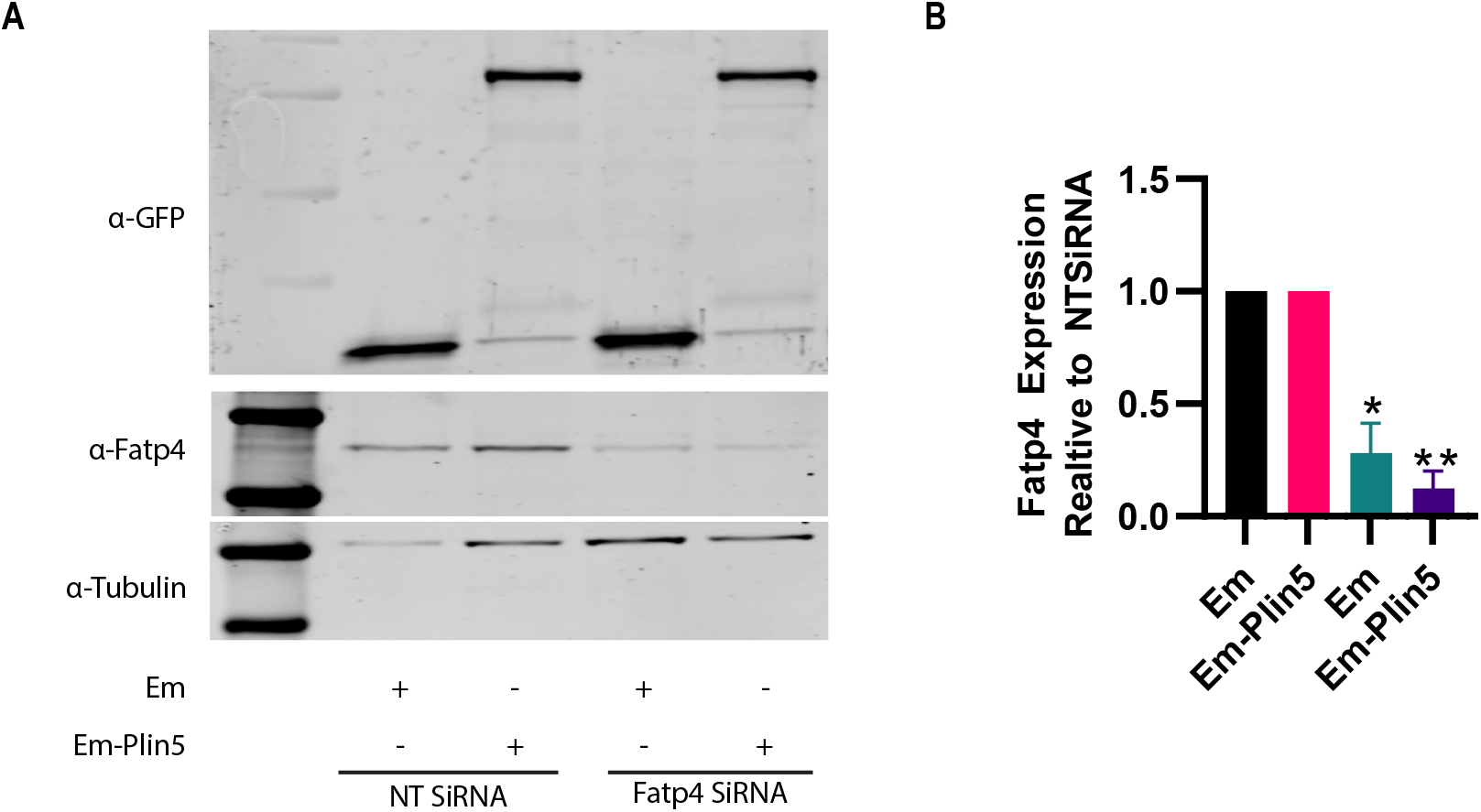
Validation of Fatp4 RNAi. **(A-B)** C2C12 myoblasts were transfected for 24 hr with non-targeting or Fatp4 siRNA. Following RNAi, cells were transfected with Em or Em-Plin5 construct. (A) Representative western blot of Fatp4 abundance following RNAi. (B) Quantitation of Fatp4 expression, normalized to nontargeting siRNA treated cells. Data are expressed as means, error bars represent ± SEM. *p < 0.05, **p < 0.01.

## Supplemental Movies

**(S1-S4)** Airyscan movies of U2OS cells transfected with Mito-RFP (magenta) and labeled for LDs with Bodipy 665/676 (green). Cells were additionally transfected with (S1) Em, (S2) Em-Plin5, (S3) Em-Plin5 (1-424) or (S4) Em-HP-Plin5 (425-463) (orange). Images were acquired every 27 s for 10 frames.

**Supplemental Table 1**

List of statistically significant Plin5-associate proteins identified in the LC-MS/MS based proteomics analysis. FC_A is enrichment relative to Em control, 2 was used as the cut off. SAINT Score is the probability a detected protein is a true interactor, 0.75 was used as the cut-off. Fold change is normalized to Plin5 abundance.

**Supplemental Table 2**

List of plasmids used in this study.

## Notes

### Competing Interest Statement

The authors have declared no competing interest.

