## Supplemental Table 1 for "Perilipin 5 interacts with Fatp4 at membrane contact sites to promote lipid droplet-to-mitochondria fatty acid transport"

| Proteins Isolated from Plin5 AP (gene name) | FC_A | SAINT Score | Fold Change: Plin5 1-443 vs WT | Fold Change: Plin5 1-424 vs WT |
|---|---|---|---|---|
| PLIN5 | 5 | 1 | 1 | 1 |
| SLC27A4 | 4.66 | 1 | 0.832509 | 0.343001 |
| FAM208A | 3.58 | 1 | 1.100372 | 0.025668 |
| ARHGEF17 | 3.23 | 1 | 0 | 0 |
| MMS19 | 3.23 | 1 | 0.715332 | 0.204618 |
| SH3BP4 | 3.03 | 1 | 0.783575 | 0.111763 |
| KIF20A | 2.85 | 1 | 0.898469 | 0 |
| ARHGEF40 | 2.8 | 1 | 1.156679 | 0.016359 |
| ZNF281 | 2.77 | 1 | 0.942393 | 0.007236 |
| ACOT9 | 2.76 | 1 | 0.715469 | 0.152222 |
| TTC37 | 2.66 | 1 | 0.43352 | 0.004746 |
| LTN1 | 2.48 | 1 | 1.753592 | 0.114778 |
| ZNF598 | 2.48 | 1 | 0.235415 | 0.080538 |
| CPNE1 | 2.4 | 1 | 0.775359 | 0.043853 |
| FKBP8 | 2.4 | 1 | 1.09269 | 0.126549 |
| SACS | 2.38 | 1 | 0.507289 | 0.046058 |
| GIT1 | 2.35 | 1 | 3.559115 | 0.031645 |
| LIG1 | 2.35 | 1 | 1.660429 | 0.035742 |
| PLCD3 | 2.35 | 1 | 1.807994 | 0.009248 |
| SYNE2 | 2.34 | 1 | 1.286989 | 0.118477 |
| PTCD3 | 2.33 | 1 | 2.617261 | 0.326183 |
| NEDD1 | 2.33 | 1 | 1.092108 | 0 |
| KIAA1217 | 2.33 | 1 | 1.407012 | 0.00984 |
| TRMT1L | 2.3 | 1 | 1.781512 | 0.015054 |
| ASAP1 | 2.3 | 1 | 1.41685 | 0.281614 |
| AURKA | 2.3 | 1 | 0.587066 | 0.454899 |
| POLRMT | 2.27 | 1 | 1.228084 | 0.189148 |
| MYO18A | 2.27 | 1 | 0.254985 | 0.037047 |
| NAE1 | 2.22 | 1 | 2.939215 | 0.161868 |
| RPAP1 | 2.2 | 1 | 2.944825 | 0.285532 |
| TMEM165 | 2.2 | 1 | 1.798197 | 0.150775 |
| DIDO1 | 2.19 | 1 | 1.005804 | 0.069992 |
| SNX2 | 2.17 | 1 | 1.54979 | 0.16127 |
| CEP131 | 2.16 | 1 | 1.404336 | 0.016073 |
| KNTC1 | 2.15 | 1 | 0 | 0 |
| PI4KA | 2.15 | 1 | 11.34677 | 0.036888 |
| EXOSC3 | 2.13 | 1 | 1.122562 | 0.030807 |
| NRBP1 | 2.1 | 1 | 1.033891 | 0.035466 |
| ZYX | 2.09 | 1 | 0.875723 | 0.218736 |

| | | | | |
|---|---|---|---|---|
| FXR2 | 2.07 | 1 | 1.52211 | 0.294445 |
| CCDC47 | 2.06 | 1 | 1.075116 | 0.16041 |
| ERBIN | 2.04 | 1 | 2.019819 | 0.213199 |
| ALDH3A2 | 2.04 | 1 | 1.270414 | 0.465539 |
| ADH5 | 2.04 | 1 | 1.135106 | 0.201224 |
| SCYL1 | 2.04 | 1 | 2.211871 | 0.555215 |
| DLG5 | 2.02 | 1 | 1.460315 | 0.130935 |
| UPF2 | 2.01 | 1 | 1.995488 | 0.022027 |
| PPIL2 | 2.01 | 1 | 0 | 0.015993 |
| DIAPH1 | 2.01 | 1 | 1.005988 | 0.069579 |
| SCAF11 | 2.01 | 1 | 0.666393 | 0.04983 |
| KIAA1671 | 2.01 | 1 | 3.388232 | 0.017262 |
| TTI1 | 2.01 | 1 | 1.126631 | 0.316291 |
| EPB41L5 | 2.01 | 1 | 0.936339 | 0 |
| NAA25 | 2.01 | 1 | 0.600513 | 0.210822 |
| CNOT10 | 2.01 | 1 | 0.985049 | 0.221627 |
| PRPS1 | 2 | 1 | 0.029489 | 0.002744 |
| RNPEP | 2.22 | 0.97 | 0.8808 | 0 |
| TERF2IP | 2.17 | 0.96 | 2.147408 | 0.038431 |
| RANBP1 | 2.2 | 0.91 | 0.498578 | 0.062687 |
| APRT | 2.04 | 0.79 | 0 | 0 |
