## Supplemental Table 2 for "Perilipin 5 interacts with Fatp4 at membrane contact sites to promote lipid droplet-to-mitochondria fatty acid transport"

| Plasmid | Source | Reference |
|---|---|---|
| mito-GFP | Jennifer Lippincott-Schwartz | |
| mito-RFP | Jennifer Lippincott-Schwartz | Mitra, K. et al. PNAS. (2009) |
| mOrange2-SKL | Michael Davidson | |
| mEmerald-C1 | Michael Davidson | addgene #53975 |
| pEYFP-C1-Plin5 | Carole Sztalryd | Wang et al. J. Lipid Res. (2011) |
| Em-Plin5 | This Study | |
| Plin5 | This Study | |
| Em-Plin5CΔ (1-424) | This Study | |
| Em-Plin5CΔ (1-443) | This Study | |
| Em-Plin5CΔ (1-453) | This Study | |
| Em-HP-Plin5C (396-463) | This Study | |
| Em-HP-Plin5C (415-463) | This Study | |
| Em-HP-Plin5C (425-463) | This Study | |
| Em-HP-Plin5C (435-463) | This Study | |
| HP-Em | This Study | |
| Pex-Em | This Study | |
| Pex-Em-Plin5C (425-463) | This Study | |
| Em-Plin5 (S155E) | This Study | |
| Em-Plin5CΔ (1-424) (S155E) | This Study | |
| Plin5 (S155E) | This Study | |
| Plin5CΔ (1-424)(S155E) | This Study | |
| Pex-Em-Plin5C (425-463) | This Study | |
| TagBFP-KDEL | Gia Voeltz | Friedman et al Science. (2011) |
| Pex-TagBFP | This Study | |
| Pex-TagBFP-Fatp4C(157-643) | This Study | |
